# Social environment affects vocal individuality in a non-learning species

**DOI:** 10.1101/2025.06.06.658121

**Authors:** Malavika Madhavan, Lucie Hornátová, Martin Šálek, Alexandra Průchová, Pavel Linhart

## Abstract

Individual recognition is fundamental to the social behaviour of many animals. In high-density populations, animals encounter and compete with conspecific rivals more frequently, which should enhance the individuality of their signals to facilitate recognition among neighbours. We investigated vocal individuality in male territorial calls of two populations of little owls (*Athene noctua*) with different densities. Further, to explore the potential influence of local population distribution on individuality, we also examined isolated males without neighbours and clumped males with neighbours. Our findings indicate higher individuality at higher densities across both scenarios, measured using two individuality metrics: Beecher’s information statistic and Discrimination score. Additionally, clumped males exhibited significantly lower acoustic niche overlaps compared to isolated males, suggesting that the immediate social environment might be more influential than larger-scale population density patterns. This study is the first to demonstrate that vocal individuality in a territorial species is influenced by conspecific density, similar to findings in group-living and colonial species.

## Background

Population density has been shown to affect many features of animal behaviour and ecology having direct and indirect effects on animals’ fitness and lifestyles. For example, population density may affect levels of intraspecific competition (Judge & De Waal, 1997; Jirotkul, 1999; Knell 2009; Cooper Jr et al., 2015), spatial behaviour such as dispersal (Luna et al., 2019) and home range size and use (Fedy & Stutchbury, 2004; Schoepf et al., 2015). High population densities may compromise fitness through declines in reproductive parameters caused by competition (Arcese & Smith, 1988) or predation (Sofaer et al., 2014). Even mating systems can be shaped by the density of a population (Emlen & Oring, 1977; Lukas & Clutton-Brock, 2013).

Population density can also shape how individuals communicate with each other (Morales et al., 2014). One of the communication tasks that might be likely affected by population density is individual recognition. Large pools of individuals and repeated interactions occurring between them may demand adaptations in the process of individual recognition, both in the design of a signal and in the receiver’s perception. For a signaller to efficiently and reliably signal their identity information, between-individual variation should consistently be much greater than within-individual variation of a signal (Beecher, 1989; Pollard et al., 2010). Strong individual signatures have been found in acoustic, visual and olfactory signals of many group-living and colonial species (Tibbetts, 2004; Charrier et al., 2010; Pitcher et al., 2010; Arnold & Wilkinson, 2011; Sheehan et al., 2016). Group-living animals represent a specific case of increased population density clusters. Animals living in larger and more complex social groups should have more complex communicative signals as suggested by the social-complexity hypothesis (Freeberg, 2006; Freeberg et al., 2012b; reviewed in Peckre et al. 2019). Studies on colonial and non-colonially breeding swallow species showed the particular importance of acoustic signal-design for individual parent-offspring recognition in species living in high population densities (Beecher et al., 1986; Loesche et al., 1991; Medvin et al.,1993). Several other studies report that species living in larger groups/colonies display larger individuality in their calls (Pollard & Blumstein 2011; Wilkinson 2003; Martin et al. 2021) and Martin et al. (2021) found that local population density affect individuality in mother-pup attraction calls in Cape fur seals (*Arctocephalus pusillus pusillus*). However, strong individual signatures are also frequently found in solitary-living, territorial species. In such a territorial context, it has been long recognized that individuals need to recognize neighbours from strangers and also among different neighbours (Brooks & Falls, 1975; Godard, 1991; Temeles, 1994; Hyman & Hughes, 2006; Jaška et al., 2015), but drivers of high individuality in territorial species were mostly neglected. Interestingly, none of the few previous studies revealed an effect of population density on vocal individuality in a territorial species (Blumstein et al., 2012; Delgado et al., 2013).

Acoustic signals are an ideal system to study identity signals because they are used for immediate long-range communication. Further, acoustic communication is highly important for nocturnal animals. Indeed, acoustic communication might have evolved to facilitate communication in reduced light conditions (Chen & Wiens, 2020). Nocturnal animals prioritise vocalisations over other channels of communication (Ramanankirahina et al., 2016; Zhang et al., 2021). Also, due to their dynamic and multidimensional nature, acoustic signals allow substantial scope for modifying signals by changing their spectral and/or temporal qualities. Therefore, nocturnal birds such as owls, are excellent models for studying acoustic communication. Owls are known to have individually distinct calls (Odom et al., 2013; Yee et al., 2016; Linhart & Šálek, 2017, see reviewed in Madhavan & Linhart, 2024) which they use to discriminate among one-another for purposes such as advertisement and territorial defence (Cavanagh & Ritchison, 1987; Grieco, 2018), mate-recognition (Nagy & Rockwell, 2012), or parent-offspring recognition (Nagy & Rockwell, 2012; Dreiss et al., 2014).

The little owl (*Athene noctua*) is a small-sized owl (body mass = 170 - 210 g) with a large natural distribution range, stretching across much of Eurasia and North Africa (Van Nieuwenhuyse et al., 2023). Little owl population densities steeply declined across different parts of its distributional range in recent decades (Van Nieuwenhuyse et al., 2023), with the most profound population declines in Central and Western Europe (Żmihorski et al., 2006; Le Gouar et al., 2011; Chrenková et al., 2017). However, in areas with favourable habitat conditions such as traditionally managed small-scale farmland with (semi-) natural grasslands (Šálek & Lövy, 2012; Mayer et al., 2021), little owls can still thrive and reach high population densities (Šálek et al., 2016; Ignatov & Popgeorgiev, 2021). As a result, little owls can be found in a wide range of population densities ranging from very low (e.g., 0.009 - 0.033 calling males / km^2^; Šálek & Schröpfer, 2008; Chrenková et al., 2017) to very high (e.g., 20 - 33 calling males / km^2^; Šálek et al. 2013; Šálek et al., 2024) with nearest neighbours being tens of metres to several kilometres apart from each other. Little owl males produce territorial ‘*hooook*’ calls that they use to defend their territories (Van Nieuwenhuyse et al., 2023). Male territorial calls have been shown to be individually vocally distinct (Linhart & Šálek, 2017), and males can also distinguish between the territorial calls of familiar neighbours and unfamiliar strangers (Hardouin et al., 2006). Further, little owls are a relatively long-lived and sedentary species, both of which contribute to stable relationships between territorial neighbours and increase the potential benefits of individual recognition (e.g. Siracusa et al 2021).

Our aim in this study was to compare the territorial calls of little owls from populations with low and high density to see if population density might drive the level of vocal individuality in a species that does not live in groups or colonies. We measured the acoustic features of little owls’ calls and used them to calculate different metrics of vocal individuality. We compared vocal individuality in two spatially separated populations with different average densities (i.e., low- and high-density populations). Further, we capitalised on the spatial distribution of the high-density population, and we compared vocal individuality in isolated males (isolated farms, with closest neighbours being several hundreds of metres apart), and clumped males living in dense aggregations of conspecifics (villages, with one to several neighbours in adjacent territories). While macroscopic differences in vocal individuality between the two populations could be explained by longer-term adaptations and genetic differences between the two populations, local differences at the level of a single population might rather suggest a plastic developmental response to current environmental conditions. We predicted the following (Figure 1):

**Figure 1.**
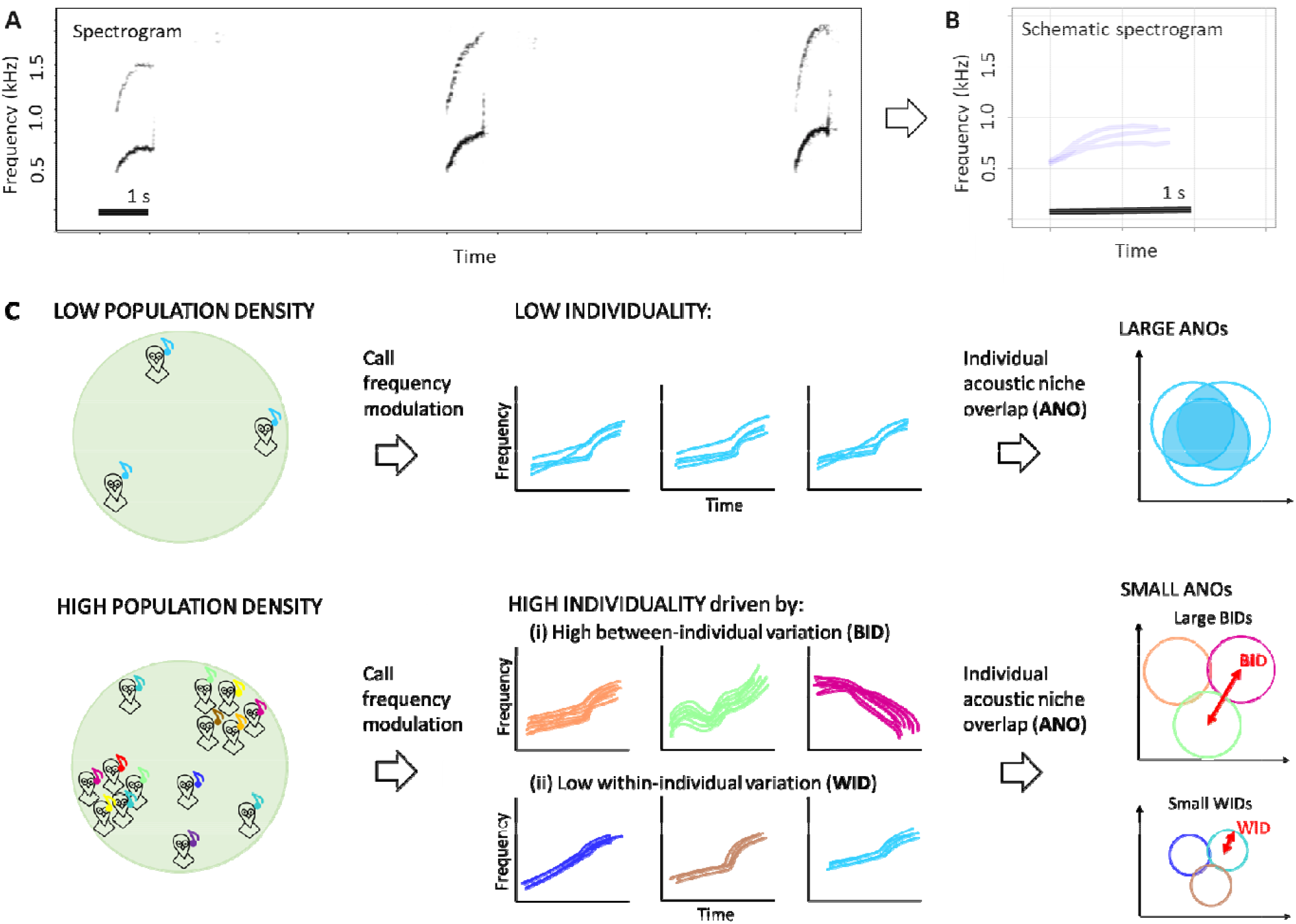
A) Spectrogram showing three territorial calls of a single little owl male and an overview of our proposed predictions: B) a schematic representation of the same three calls shown in Fig. 1A, plotted on top of each other, made using duration and peak frequency contour measurements of the calls. Fig. 1C shows a schematic representation of our predictions, showing (i) low individuality represented by large acoustic niche overlaps between individuals living in low population densities, and (ii) high individuality represented by small overlaps between individuals living in high population densities, driven by ither high between-individual variation, and/or low within-individual variation. The X and Y-axes of all plots represent time and frequency.

a. Individuals living in the high-density population have higher vocal individuality than those living in the low-density population (between population comparison).
b. Clumped individuals with neighbours will have higher individuality than isolated males (within population comparison).

Individuality has two different components: how much signals differ between individuals (between-individual variation) and how consistent they are within each individual (within-individual variation) (Falls, 1982; Beecher, 1989; Pollard et al., 2010). Increase in individuality can be mechanistically driven by an additional increase in between-individual variation or by reducing within-individual variation, or both. Therefore, we characterised each of these individuality components separately and compared their values depending on population density, to explain the source of any possible change in individuality related to population density.

## Methods

### Recordings

Little owl males were recorded between March and April in both 2013 and 2014. This period overlapped with the courtship period, when little owl males most actively produce territorial hoots (Exo, 1988). The research was conducted in two study sites: the first one in Northern Bohemia, Czech Republic (hereafter ‘LOW density population’) (GPS: 50.41°N, 14.04°E), and the second close to Hortobágy National Park, Hungary (hereafter ‘HIGH density population’) (GPS: 47.57°N, 20.93°E). The average little owl density in the LOW-density population was 0.09 calling males per 10 km^2^ (Šálek & Schröpfer, 2008), and in the HIGH-density population, the average density was 5.01 calling males per 10 km^2^, which is one of the highest reported densities in central Europe (Šálek et al., 2013). However, within the human settlements of the Hungarian population, densities of little owls are even higher and can reach up to 33.3 calling males per km^2^ (Šálek et al., 2013, Šálek et al., 2024), males therefore frequently have one or several calling neighbours within a 200 m radius (hereafter ‘CLUMPED males’ - high local population density). On the other hand, isolated males without any detected neighbours were found on isolated farms in the same population (Šálek et al., 2013). The closest human settlements to these isolated farms, which might provide suitable breeding habitats for little owls, were 552 - 1445 m away (Průchová et al., 2024). The farms being surrounded by farmland or grassland had no other nesting opportunities for little owls (hereafter ‘ISOLATED males’ - low local population density), as breeding sites of little owls are currently exclusively located in human-made structures (Šálek et al., 2013; Šálek, 2014; Chrenková et al., 2017). Presence of neighbours was noted during initial acoustic surveys and confirmed by scanning the recordings for territorial calls of the focal male and any other males (for more details on survey methods, see Šálek et al., 2013). Territorial calls of male little owls were recorded in optimal weather conditions (no strong winds or rain), and during the peak of calling activity (between dusk and midnight). For comparison between populations (HIGH vs. LOW-density populations), recordings were made with a Marantz PMD660 solid-state recorder (sampling frequency: 44,100 Hz, 16-bit, WAV format without any compression) and a Sennheiser ME67 directional microphone with Rycote windshield protection. Males were recorded after ca. 1-minute-long playback provocation, from up to 50 m from the focal individual. For the comparison within the HIGH-density population (CLUMPED vs. ISOLATED males) spontaneous calls without playback provocation were used. Recordings were made using autonomous recorders Olympus DM-650 programmed to record continuously over night from 7 PM to 7 AM (sampling frequency: 44 100 Hz, 16-bit, WMA format). Recorders were positioned within 50 metres from the preferred calling post identified during the previous population census with the use of playback. Often, recorders operated on the localities over two consecutive nights. Nights containing more hooting sequences of better quality were selected for further analysis. Overall, 35 recordings were made in 2013 (13 in LOW; 11 in HIGH; 6 in CLUMPED, 5 in ISOLATED) and 33 recordings were made in 2014 (9 in LOW; 11 in HIGH; 8 in CLUMPED; 5 in ISOLATED). No male was recorded in both years, and very few males were recorded with both playback and spontaneous methods (see Supporting information, Figure S1).

### Processing of acoustic data

All processing of recordings, and measurements were done in RavenPro version 1.6 (Charif et al., 2010). As the fundamental frequency of calls was always between 500 Hz and 2000 Hz (mean ± standard deviation of minimum frequency = 632 ± 117; maximum frequency = 1003 ± 109), we band pass filtered recordings (500 Hz - 2000 Hz) and reduced the sampling frequency to 4000 Hz before analyses. Spectrograms of all recordings were produced using the following settings: window size = 512 samples, with Hann type window, window overlap = 93.75% (temporal resolution = 8 ms/32 samples; spectral resolution = 7.81 Hz).

In the first analysis (comparing individuality in HIGH- and LOW-density populations), we used recordings obtained from 22 territorial males for each population (29 ± 9 calls per male in LOW and 26 ± 9 calls per male in HIGH population). In the second analysis (comparing individuality in CLUMPED and ISOLATED males), we used spontaneously recorded calls from 24 males. This consisted of 14 CLUMPED males and 10 ISOLATED males. 25 ± 1 calls from CLUMPED males, and 24 ± 1 calls from ISOLATED males, with minimal background noise from each male were selected and measured from sequences occurring at different times throughout the night after evaluating for their quality (no overlap with background noise, wind., etc.).

For both analyses, we measured the duration and frequency modulation (Peak Frequency Contour - PFC in RavenPro) of calls. Frequency modulation is very prominent in little owl male calls. It has been shown to be individually distinct and perform better for individual identification than using spectral features or spectrogram cross-correlation (Linhart & Šálek 2017). PFC measurements were visualised and manually reviewed. Calls in which the contour was not properly traced were omitted from further analyses. As RavenPro takes PFC values for each spectrogram slice of the call, the number of PFC values varies depending on the duration of the call, however, further analyses required the same number of variables for each call. To overcome this problem, we used the R package *Rraven* (Araya-Salas, 2020) to conduct dynamic time warping on the calls, using the function ‘*extract_ts*’. This allowed us to extract 10 PFC measurements for each call, which are spread out evenly through the call even when call durations differed. Previously, 10 sampling points were found to be adequately representative of the PFC for individual identification (Linhart & Šálek 2017).

### Statistical analyses

#### Quantification of vocal individuality

We used different metrics to assess individuality in little owl territorial calls. Beecher’s statistic (HS), and Discrimination score (DS) are two common sample-wide individuality metrics (i.e. a single individuality value is obtained for the whole population of individuals). DS has been used widely in different studies, but HS is better for comparison between studies with different sampling (Beecher et al., 1989; Linhart et al., 2019). HS and DS were both calculated using the R package *IDmeasurer* (Linhart et al., 2019). For statistical assessment of individuality, we used metrics obtained at the level of each individual: average within-individual variation and average between-individual variation were assessed to investigate changes in each of the individuality components separately, and an aggregate acoustic niche overlap was calculated as a metric integrating both components together.

#### Beecher’s information statistic (HS)

is a method rooted in information theory, which quantifies how well a trait can function as a signal of individual identity (Beecher, 1989). It is calculated using the ratio of total variation (total sum of squares) of the specific condition (HIGH, LOW, CLUMPED, ISOLATED) to within-individual variation (within-sample sum of squares). It lies on a scale from zero to infinity, with zero meaning no individuality. The higher the value of HS, the easier it is to discriminate between individuals. HS has important advantages over the other metrics (Linhart et al., 2019) and can serve as a comparative metric across species (Medvin et al., 1993; Linhart et al., 2019). It has been used to calculate individuality in several studies, across species and sensory modalities (Blumstein & Munos, 2005; Pollard & Blumstein, 2011; Linhart & Šálek, 2017; Tumulty et al., 2022). As HS requires the input trait variables to be uncorrelated, we first centred and scaled all variables (10 PFC values and duration) and then performed a Principal Components Analysis (PCA) on them using the function ‘calcPCA’ (package *IDmeasurer*). The function ‘calcHS’ from the same package calculates HS for each principal component and then sums them through the whole call to compute the overall Beecher’s statistic HS of the signal. The PCA plots can be found in the Supporting information (Figures S2 – S3).

#### Discrimination score (DS)

is the accuracy score of calls being correctly classified to an individual and is expressed as the percentage of correctly classified calls. We used the function ‘calcDS’ (package *IDmeasurer*), which uses Linear Discriminant Analysis (LDA) with leave-one-out cross-validation to compute the DS for the whole pool of individuals. LDA has been widely used in studies reporting vocal individuality (Galeotti et al., 1993; Favaro et al., 2015; Li et al., 2017). However, it is more susceptible to estimation biases caused by the number of included individuals and calls per individual. To prevent these biases, we aimed to select similar numbers of individuals and calls for our comparisons.

#### Within and between individual variation (WID, BID)

To be an efficient means of recognition at the individual level, the between-individual variation of an identity trait must be consistently greater than the within-individual variation of the same trait (Beecher, 1989; Pollard et al., 2010). We used the PCA scores from previous analyses and Euclidean distances among the calls to characterise within and between-individual variation. To assess within-individual variation, we measured and averaged the Euclidean distance from each call of an individual to the individual’s centre (mean value of all individual’s calls, WID). High individual call consistency would be characterised with low WID values. To assess between-individual variation, we measured the Euclidean distances from the centre value of each population to the centre value of each individual of that population calculated as described above (BID). Larger BIDs would represent an increase in acoustic space occupied by individuals within each population.

#### Acoustic niche overlap (ANO)

Niche theory emerged as describing each species in an ecosystem to have an ecological niche which occupies ‘n-dimensional hypervolumes’ (Hutchinson, 1957). Ecological niches and their overlaps are typically used to study resource partitioning between different species in an ecosystem. Since then, there have been developments in how niches are calculated as well as its applications in different branches of ecology (Bearhop et al., 2004; Raxworthy et al., 2007; Alvarado□Serrano & Knowles, 2014). In animal communication, the concept of ecological niche has been applied to studying separation of acoustic space between species (Henry & Wells, 2010; Sinsch et al., 2012; Hart et al., 2020). A similar logic can be also applied to the partitioning of acoustic space among different individuals belonging to the same species. We assume that high individuality results in low overlaps of acoustic features between different individuals, enabling reliable discrimination among individuals. The R package *‘nicheROVER’* (Swanson et al., 2015) was designed to calculate multidimensional ecological niche region sizes and the overlaps of those niches; we adapted this approach to calculate individual acoustic niches. The package ‘nicheROVER’ (function: ‘overlap’) finds the probability of any call produced by the focal individual being within the acoustic niche of another individual. We calculated all pairwise niche overlaps within each sample of individuals (alpha = 99%, number of Monte Carlo draws = 10,000). Then, we calculated the final aggregate acoustic niche overlap metrics for each individual by summing the probabilities of that individual being in the acoustic niche of any other individual of its population.

### Comparing populations and subpopulations

HS and DS values could not be statistically compared as they represent population-wide metrics. We used non-parametric Wilcoxon’s rank sum tests (R package *‘stats’* version 4.2.0) to compare WIDs, BIDs, and ANOs between the different population density conditions (HIGH vs. LOW and CLUMPED vs. ISOLATED). The sample size for different statistical tests was always the same (HIGH: n = 22; LOW: n = 22; CLUMPED: n = 14; ISOLATED: n = 10). All statistical analyses were carried out in R version 4.2.0 (R Core Team, 2022). We used the value of α = 0.05 as a threshold for significance of the results.

## Ethical Note

Recordings were made in places with unrestricted public access and on wild animals. This study was purely observational and non-invasive; therefore, no special permits were required.

## Results

### Individuality in HIGH- and LOW-density populations

We calculated Beecher’s statistic and discrimination score, and both metrics indicated higher individuality in the HIGH-density population (HS = 6.38, 83 unique signatures; DS = 87.7%, 22 individuals, 26 ± 9 calls per individual) compared to the LOW-density population (HS = 4.96, 31 unique signatures; DS = 79.4%, 22 individuals, 29 ± 9 calls per individual). We did not find a significant difference in either WID values (*W = 198; p = 0*.*31, HIGH density, median = 0*.*88, Inter-quartile range (IQR) = 0*.*42; LOW density, median = 1*.*12, IQR = 0*.*84)* or BID values (*W = 262; p = 0*.*65, HIGH density, median = 2*.*81, IQR = 1*.*22; LOW density, median = 2*.*41, IQR = 2*.*08*) between both populations. Despite not seeing significant differences between the two groups, we still observe that the median values for both comparisons are in the direction of higher individuality in the HIGH-density population. Similarly, when comparing the ANOs of male calls in HIGH and LOW density populations, we did not find a significant difference, but observed that ANOs in the HIGH density population tended towards being lower than those in the LOW density population (*W = 165, p = 0*.*07, HIGH density, median = 0*.*0059, IQR = 0*.*05; LOW density, median = 0*.*0357, IQR = 0*.*16;* Fig. 2a).

**Figure 2.**
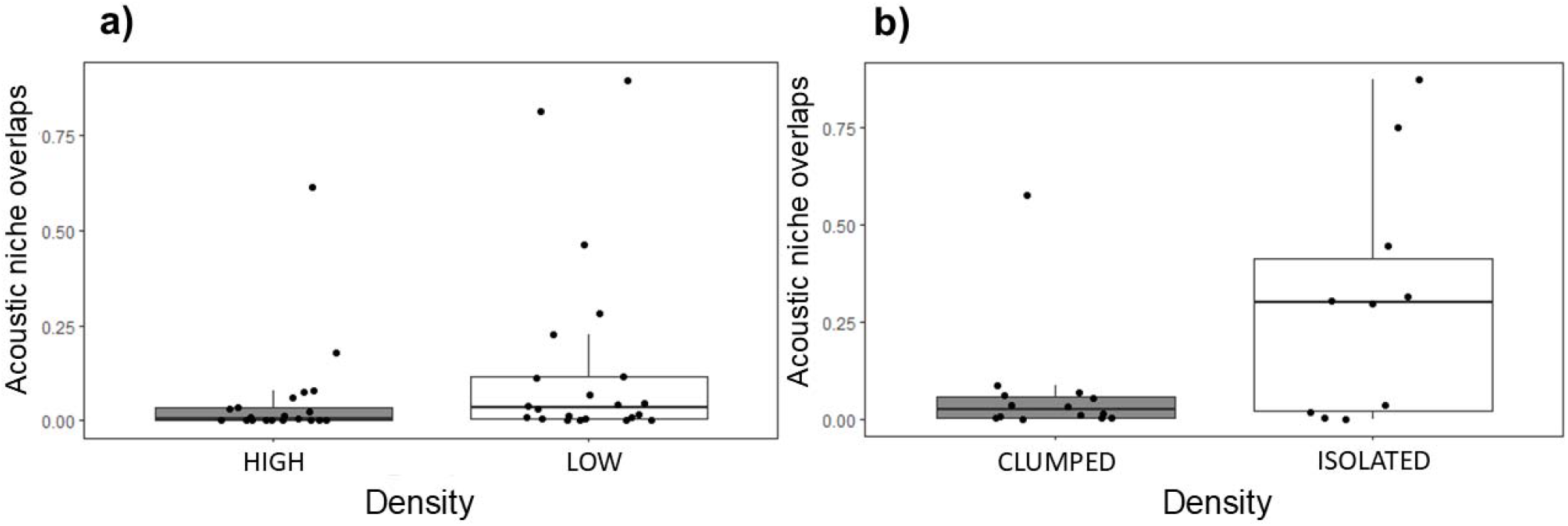
Boxplot with differences in aggregate ANO values **a)** Between populations; HIGH density and Low (LOW density) and **b)** Within population; High (CLUMPED males with neighbours) and Low (ISOLATED males without neighbours). Medians, quartiles and non-outlier minima and maxima, along with original data points (dots) are displayed.

### Individuality in CLUMPED and ISOLATED males

As in the first analysis, both metrics indicated higher individuality in the high local population density males (CLUMPED males; HS = 5.29, 39 unique signatures; DS = 87.5%, 14 individuals, 25 calls per individual) when compared to the low local population density males without neighbours (ISOLATED males; HS = 3.98, 15 unique signatures; DS = 78.9%, 10 individuals, 25 calls per individual). Here again, we did not find a significant difference in either WID values (*W = 61; p = 0*.*63, CLUMPED males, median = 1*.*31, IQR = 0*.*47; ISOLATED males, median = 1*.*22, IQR = 1*.*03)* or BID values (*W = 81; p = 0*.*55, CLUMPED males, median = 2*.*35, IQR = 1*.*45; ISOLATED males, median = 2*.*32, IQR = 0*.*56*) between the two density conditions. However, aggregate ANOs in CLUMPED males was significantly lower than those of ISOLATED males (*W = 33, p = 0*.*04, CLUMPED males, median = 0*.*0223, IQR = 0*.*05; ISOLATED males, median = 0*.*2997, IQR = 0*.*39;* Fig. 2b), suggesting a higher level of individuality in CLUMPED males.

## Discussion

Here, for the first time, we document that vocal individuality is affected by the population density in a species that does not live in social groups or colonies. This is supported by two population-wide individuality metrics. Further, we used additional metrics that allow statistical comparisons of individuality between populations. We did not observe significant differences in individual acoustic niche overlaps between the two populations with contrasting densities. However, within the high-density population, local aggregation of males (CLUMPED males) had significantly smaller acoustic niche overlaps (i.e., higher individuality) than ISOLATED males. When investigating potential mechanisms leading towards higher vocal individuality in CLUMPED males, we did not find significant support for CLUMPED males making their calls more distinct from other males (larger between individual distances, BID), or for them calling in a more consistent way (smaller within individual distances, WID). Higher individuality in CLUMPED males thus seems to be a result of a smaller cumulative combination of both mechanisms.

Our findings are broadly consistent with the social complexity hypothesis (Freeberg, 2006), where increased complexity in acoustic signalling - including individuality, have been linked with increasing social complexity across many species (see Peckre et al., 2019 for a detailed account of such examples). High population density is considered one among the different proposed indicators of increasing social complexity. However, studies dealing with the social complexity hypothesis mostly focus on species’ vocal repertoire rather than individually distinct vocalisations of the same call-type. The few studies that address communication complexity in individually specific calls, typically study these mechanisms at an interspecific level (but see Martin et al. 2021), and in group-living or colonial species. Higher individuality has been found in animal species living in larger or more complex groups (Wilkinson, 2003; Pollard & Blumstein, 2011; Charrier 2020). Perhaps one of the best examples so far of individuality being selected for at high densities is shown in the comparative study of four swallow species (Beecher et al., 1986). Chicks of sand martin (*Riparia riparia*) and northern rough-winged swallow (*Stelgidopteryx serripennis*), which are colonial breeders, show much higher vocal individuality than those of the closely related barn swallows (*Hirundo rustica*) and cliff swallow (*Hirundo pyrrhonota*), which are non-colonial breeders.

Very few studies compared individuality between populations of different densities, in a single species. Previous work on Cape fur seals found that the individuality in two call-types involved in parent-offspring communication was higher in the larger colony (Martin et al., 2021). The authors also found high individuality in the territorial calls of male Cape fur seals; however, they did not have data to compare individuality in territorial calls in both populations. They use the index of vocal stereotypy (IVS) as an individuality metric in their study which corrects DS by chance levels (IVS = DS / chance assignment; chance assignment = 100 / number of individuals). This metric has the same issue as DS, as also discussed by the authors, i.e. it depends heavily on the number of individuals in the sample. Recalculating our values indicates that the IVS values and differences between studied conditions in our study would be comparable if not exceeding those found by Martin et al. (2021, Figure 2) for comparable number of individuals (IVS HIGH = 19.29; IVS LOW = 17.47; IVS CUMPED = 12.25; IVS ISOLATED = 7.89).

To our knowledge, our study is the first work to show that high population density is associated with increase of vocal individuality in territorial calls of a monogamous, pair-living species. Our results contradict two previous studies on the same topic. Blumstein et al. (2012) studied individuality in seven species of passerines and did not find any relationship between their population density and vocal individuality in its territorial songs. This study analysed features of entire songs coming from different species with quite varied vocalisations. However, songbirds may encode individuality specifically in certain parts of their songs (Elfström, 1990; Wegrzyn et al., 2009). Therefore, the analysis might be confounded by including song parts with different signalling functions. In the second study, Delgado et al. (2013) suggested that high population density may even reduce vocal individuality in Eurasian eagle-owls (*Bubo bubo*). However, they analysed vocalisations within a single population only and compared their results with those of a different study (Lengagne, 2001). However, the results of both studies are not directly comparable (slightly different sets of extracted acoustic features, different methods of feature extraction, different spectrogram settings, different sample sizes, etc.). Actually, both studies indicate very high vocal individuality with the difference in DS smaller than in our study (Delgado et al., DS = 95.8%; Lengagne et al., DS = 100%). Our study overcomes both these previous shortcomings by focusing on simple calls that function in individual recognition (Hardouin et al., 2006) and by direct comparison across populations using the same analysis methods.

Our study does not come without limitations. We realise that comparing two populations can lead to false generalisations, because differences between two populations can arise due to many different reasons besides the population density. Ideally, we would need to have a sufficient number of replicates from other HIGH-LOW density and CLUMPED-ISOLATED conditions. On the other hand, this is a problem of many other pioneering studies. We believe our results could be still highly useful for future studies and comparisons. While we could not include more replicates, we used two different approaches (high-low density, clumped-isolated males) and came to very similar results in both cases. The comparison between the two approaches also suggests that focusing on the immediate social environment is probably more promising for future studies.

Further, higher vocal individuality in the Hungarian population was indicated by higher discrimination score and Beecher’s information statistic, which are both population-wide metrics. However, this pattern was not confirmed by significant differences in individual acoustic niche overlaps like in the case of comparison between CLUMPED and ISOLATED males. One reason possibly hindering the assumed difference in acoustic niche overlaps between populations might be the spatial distribution of males in both populations. HIGH and LOW populations indeed both included some ISOLATED and some CLUMPED males which could diminish possible differences between the populations if local spatial distribution is the decisive factor.

Another explanation could lie in the fact that recordings were collected by different methods for both analyses. For the first comparison between populations, recordings were made after playback provocation, while recordings of spontaneous calls were used to compare individuality in CLUMPED and ISOLATED males within the high-density population. The use of playback has been shown to have effects on vocal responses of different species including changes in vocal output, timing, frequency and content (McGregor et al., 1992; Hardouin et al. 2007; Linhart et al. 2013; Geberzahn & Aubin 2014; Thomsen et al. 2019). Therefore, use of playback, in general, might be assumingly associated with increased within-individual variation of calls eventually leading to smaller differences in the case of HIGH and LOW conditions. On the other hand, studies on owls successfully use playback elicited calls to identify individuals within and across breeding season, suggesting little effect of playback on individually distinct call features (Delport et al., 2002; Tripp & Otter, 2006; Rognan et al., 2009; Takagi, 2020). We cannot provide direct assessment of effects of playback on vocal individuality as we currently do not have enough good quality recordings for a proper analysis. However, consistent PFCs patterns can be found in spontaneous and provoked recordings of the same males (Supporting information, Figure S1). On the other hand, we also observed that PFC can change during and across years in both playback-provoked as well as in spontaneous calls, suggesting that PFC can be consistent or substantially modified in both spontaneous as well as in provoked recordings (PL, unpublished results).

Various mechanisms exist that could be responsible for the higher vocal individuality in CLUMPED males compared to males living on isolated territories. For example, neighbours with dissimilar calls might settle next to each other leading to the establishment of vocally diverse clusters. Sexual selection for high consistency of calls observed in some species (Rivera-Gutierrez et al., 2011; Sierro et al., 2023) might accidentally result in higher individuality. However, we did not observe a significant decrease in within-individual variation in our study that should be associated with higher consistency of calling. Further, individuals in urban habitats (most of our CLUMPED males came from villages and ISOLATED males from rural areas) might be more diverse genetically, and consequently acoustically. However, a review on this topic indicates rather lower genetic diversity in urban populations (Miles et al., 2019). In following studies, genetic samples might allow us to take individual relationships and genetic diversity of populations into account to rule out the possibility that higher vocal individuality is merely a consequence of larger genetic divergence within a population / sub-population.

Our study might also represent an example of some form of vocal plasticity in owls. Indeed, calling of little owls seems quite plastic, and this fact is reflected in descriptions of their vocal repertoire (Van Nieuwenhuyse et al., 2023), but owls are typically considered to be vocal non-learners (Brenowitz, 1991; Ten Cate, 2021). Simple forms of vocal plasticity include short-time modifications of call amplitude, pitch and timing in response to noisy conditions or conspecific calls; these are taxonomically widespread (Brumm & Zollinger, 2011; Dunlop et al., 2014; Halfwerk et al., 2016) but have rarely been described in owls. Moreover, little owls would need to modify both the duration and frequency modulation of their calls to diverge from their neighbours. Individuals could also have several variants of calls in their repertoire and might use a dissimilar call in interactions with their neighbours to enhance individual recognition among local neighbours - a form of a contextual vocal learning (Janik & Slater 2000). At the extreme, our study might represent an example of innovation learning (Janik & Slater, 2000) where calls of animals diverge from the models. Observations documenting long-term divergence in calls, consistent with innovation learning, were described in bottlenose dolphins (*Tursiops truncatus*), where whistle signatures of daughters were more dissimilar from their mothers than whistle signatures of sons (Sayigh et al., 1995); in common loon males (*Gavia immer*), who changed the structure of their yodels to sound more different than those of the previous territory owner (Walcott et al., 2006); in green-rumped parrotlets (*Forpus passerinus*), where nestlings in larger broods had higher diversity in contact calls than those in smaller broods (Arellano et al., 2022). Call divergence between individuals through vocal learning is also among the suspected mechanisms behind the fall of dialects in male elephant seals (*Mirounga angustirostris*) (Casey et al. 2018). While there is some evidence that bird clades previously thought to be vocal non-learners actually possess abilities that fall squarely within the boundaries of what constitutes vocal production-learning (Ten Cate, 2021), fascinating speculations about any form of vocal learning in owls would need to be confirmed by additional research.

## Conclusion

In this study, we show that vocal individuality in the territorial calls of little owls is influenced by their social environment, particularly, by the population density. Similar results were previously reported only for group-living and colonial species. While qualitatively similar patterns were found at both spatial scales, our results indicate that distribution within local neighbourhoods (ISOLATED vs. CLUMPED males) is more important than larger-scale population characteristics (HIGH vs. LOW average population density). Limitations of our study do not allow us to conclusively decide about the mechanism leading to patterns observed here. In the future, data is needed to also take into account the spatial and genetic relationships between individuals. Further, more longitudinal studies would be needed to document the development of individually distinct calls in owls, and their plasticity later in life.

