## Supporting figure S1 for "Social environment affects vocal individuality in a non-learning species"

**A)**

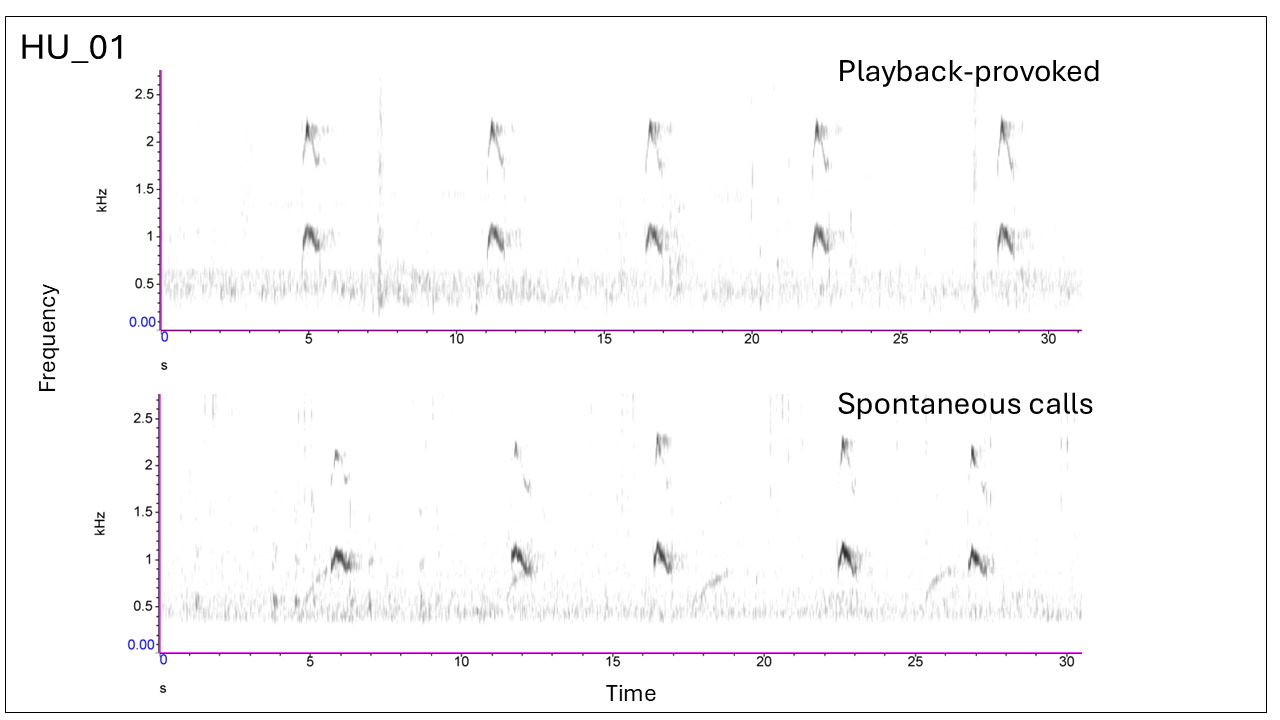

**B)**

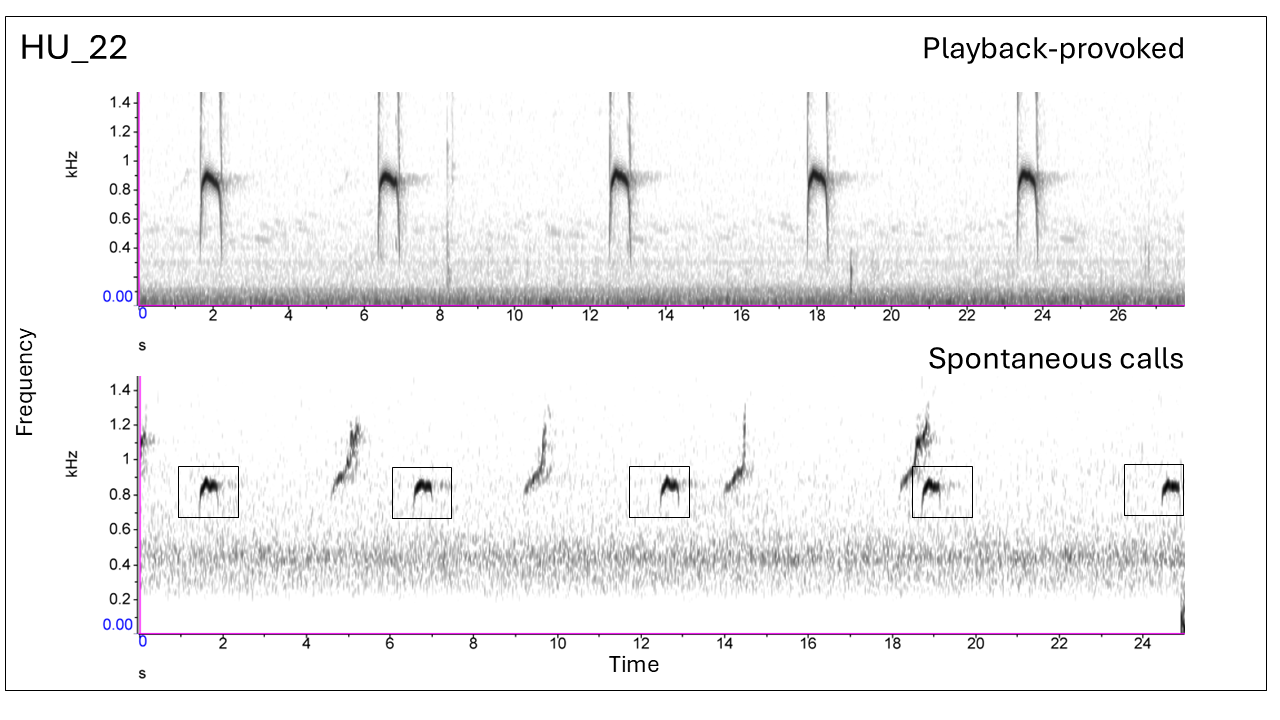

**Fig. S1.** A) and B) show spectrograms of calls recorded using playback provocation and spontaneous calls recorded using passive acoustic recorders, for individuals HU_01 and HU_22 respectively.
