## Supporting figures S2-S3 for "Social environment affects vocal individuality in a non-learning species"

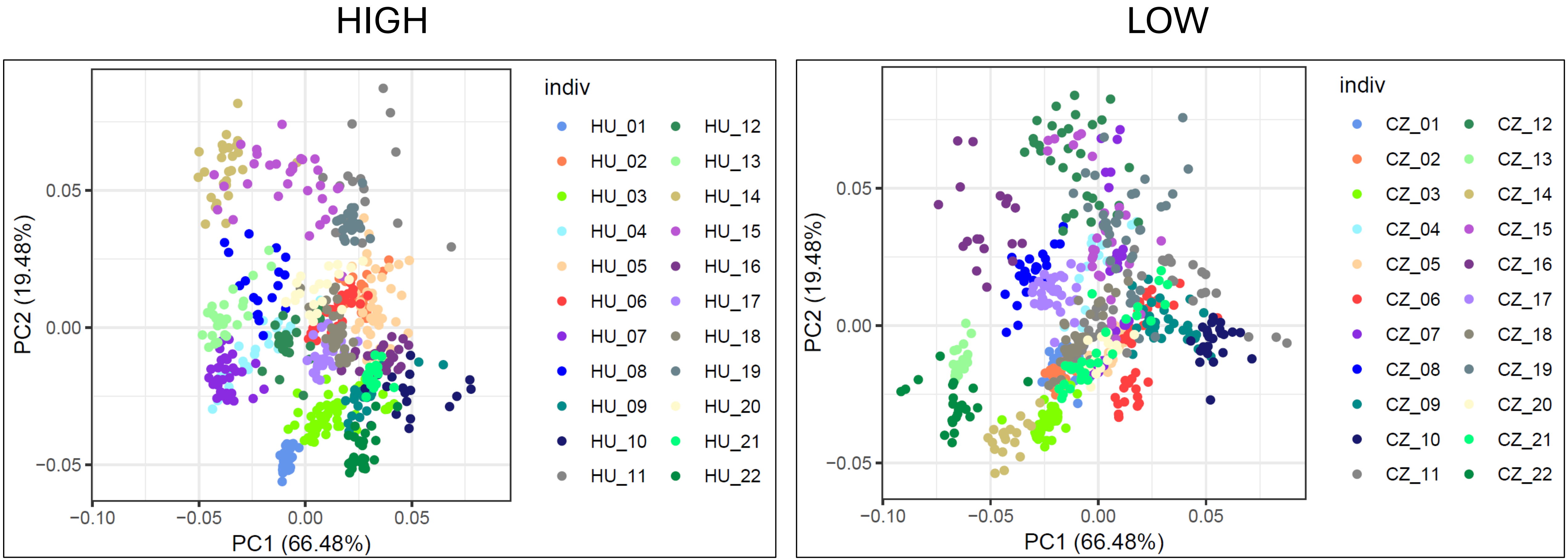


**Fig. S2.** Plots from a Principal Components Analysis done on call components (10 peak-frequency points and duration) for males from both HIGH and LOW density conditions.

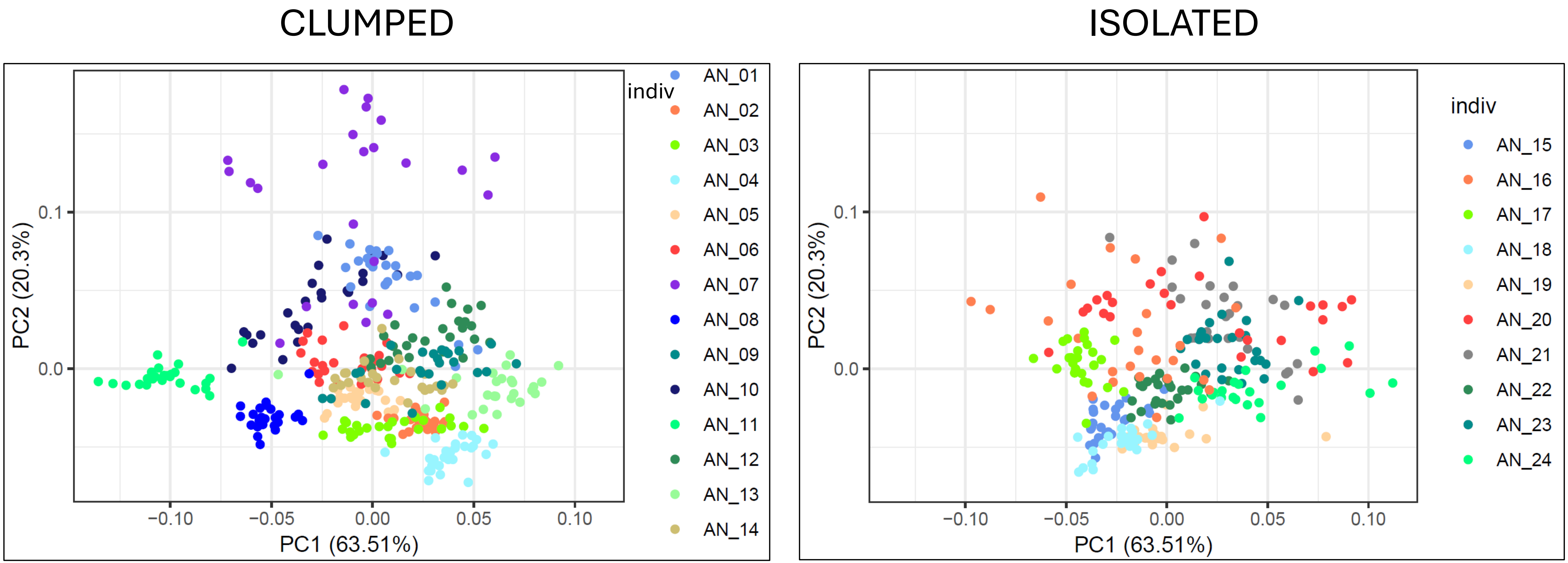


**Fig. S3.** Plots from a Principal Components Analysis done on call components (10 peak-frequency points and duration) for both CLUMPED and ISOLATED males.
